# The effect of stress on memory for temporal context

**DOI:** 10.1101/2021.04.23.441105

**Authors:** Nicole D. Montijn, Lotte Gerritsen, Iris. M. Engelhard

**Affiliations:** Department of Clinical Psychology, Utrecht University, The Netherlands

**Keywords:** Stress, Episodic Memory, Temporal Context, Time Perception, Learning

## Abstract

Stress and emotional arousal interfere with encoding of temporal context memory for episodic events. However, it remains unclear how stress affects more fine-grained temporal memory, such as episodic events sequences and event times. Here, 86 healthy participants (M age = 22.5; 46% women, 54% men) were subjected to either a stress condition (socially evaluated cold pressor test) or a control condition, directly after or at a delay of 30 minutes they were presented the temporal structure of four virtual days. In these virtual days, time was scaled and participants could use clock cues to construe the passage of time within a day. We examined whether acute stress would interfere with encoding of episodic event sequences and temporal memory. Our results show that when learning took place directly after a stressor, virtual time estimates were more strongly biased towards a generalized timeline but temporal memory overall was not differentially affected between the stress and control groups. Exploratory analyses suggest that memory accuracy improved in men and deteriorated in women as a function of subjective stress levels following acute stress. In conclusion, acute stress amplified memory generalization but we found no stress related differences in memory accuracy across levels of temporal granularity,.

## Introduction

Stress and arousal influence how well we remember events. Studies show that they typically enhance recall for emotional information at the detriment of contextual information, such as temporal context memory (Huntjens et al., 2015). These findings are generally in line with insights from stress researchhereafter SECPT; (Ehlers & Todd, 2017; Sagliano et al., 2018; Schwabe, Bohringer, et al., 2008) that show selective attention and memory for material that is directly related to the stressor and thus might promote survival.

Interestingly, the same memory enhancing benefits that are typically imbued on emotional material also occur for neutral events (Clewett & McClay, 2021; Tambini et al., 2017). For instance, Tambini et al. (2017) showed that exposure to several blocks of emotional stimuli (IAPS pictures) induced a prolonged state of arousal that enhanced memory for neutral stimuli shown 9 to 33 minutes later. However, the same task was used for the emotional and neutral blocks, so it is unclear to what extent temporal context memory was also affected. A recent study that examined the effects of arousal on temporal order memory for neutral item pairs showed that sequence memory was enhanced for item pairs that followed an unrelated arousing stimulus, but it was impaired when the arousing stimulus separated the item pair (Clewett & McClay, 2021). The latter effect may be caused by contextual-shift that is induced by the sudden onset of an emotional stimulus which induces an event boundary that separates the emotional experience from what came before (Clewett et al., 2019). In contrast, as shown by Tambini et al. (2017) neutral information that follows an emotional event boundary may be unjustly lumped into the emotional experience.

The work we discussed thus far gives some insight into the likely fate of memories encoded during emotional arousal. Yet is remains unclear to what extent emotional arousal affects temporal granularity on a finer scale, and whether the effects remain over a longer time span (e.g., one day later). Temporal context captures when an episodic event occurred in relation to other events, such as event order and time distance between events, thereby providing a framework by which we can mentally organize and cluster events. Accurate retention of this temporal event context ensures that information is interpreted within the appropriate contextual boundaries, and it protects against overgeneralization of memory (Moore & Zoellner, 2007; Wang et al., 2022), and presumably the development of intrusive memories (e.g., Al Abed et al., 2020; Brewin, 2014).

Here, we investigated the time-dependent effect of stress on encoding the temporal structure of episodic event sequences using a 2-day paradigm. Participants were trained to learn the temporal structure of four virtual days (Bellmund et al., 2022) either directly after stress or control task or after a 30-minute delay (wait and no wait condition). The wait manipulation allowed us to look at time dependent effects of stress on memory due to the direct release of noradrenaline and delayed glucocorticoid response (Joëls et al., 2011; Schwabe & Wolf, 2014; van Ast et al., 2013). The next day, we assessed memory for the temporal structure of the virtual days. We expected that the control groups would retain a detailed representation of temporal structure, as was found in a previous study using this task (Bellmund et al., 2022), while the stress groups would retain temporal information at a higher level of granularity.

As a secondary objective, we sought to investigate whether possible stress-related reductions in temporal memory accuracy were directional. A previous study using the same temporal learning task found that individual event time estimates tended to be biased towards the average time of events in the same sequence position (Bellmund et al., 2022). This type of generalization bias was present in participants with relatively accurate temporal memory, but it appeared to be enhanced when memory specificity was low. If arousal reduces specificity of temporal memory, then this generalization bias should be more prominent in the stress groups compared to controls.

## Methods

### Participants

Participants were 86 cis-gender adults (40 female, 46 male, M age = 22.5, range = 18 – 32) with no self-reported current psychiatric disorders. They were recruited on campus using flyers, as well as through social media. Female participants were required to be on hormonal birth control to control for potential bias by fluctuations of female hormones (Espin et al., 2013). In the Netherlands, this is the most widely used form of contraceptives, especially amongst students (source CBS: https://www.cbs.nl/nl-nl/nieuws/2014/25/gebruik-pil-daalt-spiraaltje-wint-terrein). Participants were randomly assigned to one of four experimental groups; stress no wait, stress 30-min wait, control no wait, control 30-min wait (Table 1). All participants provided written informed consent. A power analysis (G*Power Version 3.1; Faul, Erdfelder, Buchner, & Lang, 2009) based on prior research (Schwabe & Wolf, 2010) showed that a sample size of at least 20 per group was necessary to detect an effect of stress and timing on memory encoding (power = .90, η*_p_*^2^ = 0,12). Taking potential missing values into account, we aimed to test 20-25 participants per group. They were remunerated with course credit or had the chance to win a gift card for their participation. The study was approved by the institutional ethical review board at Utrecht University (FETC16- 090).

**Table 1.**
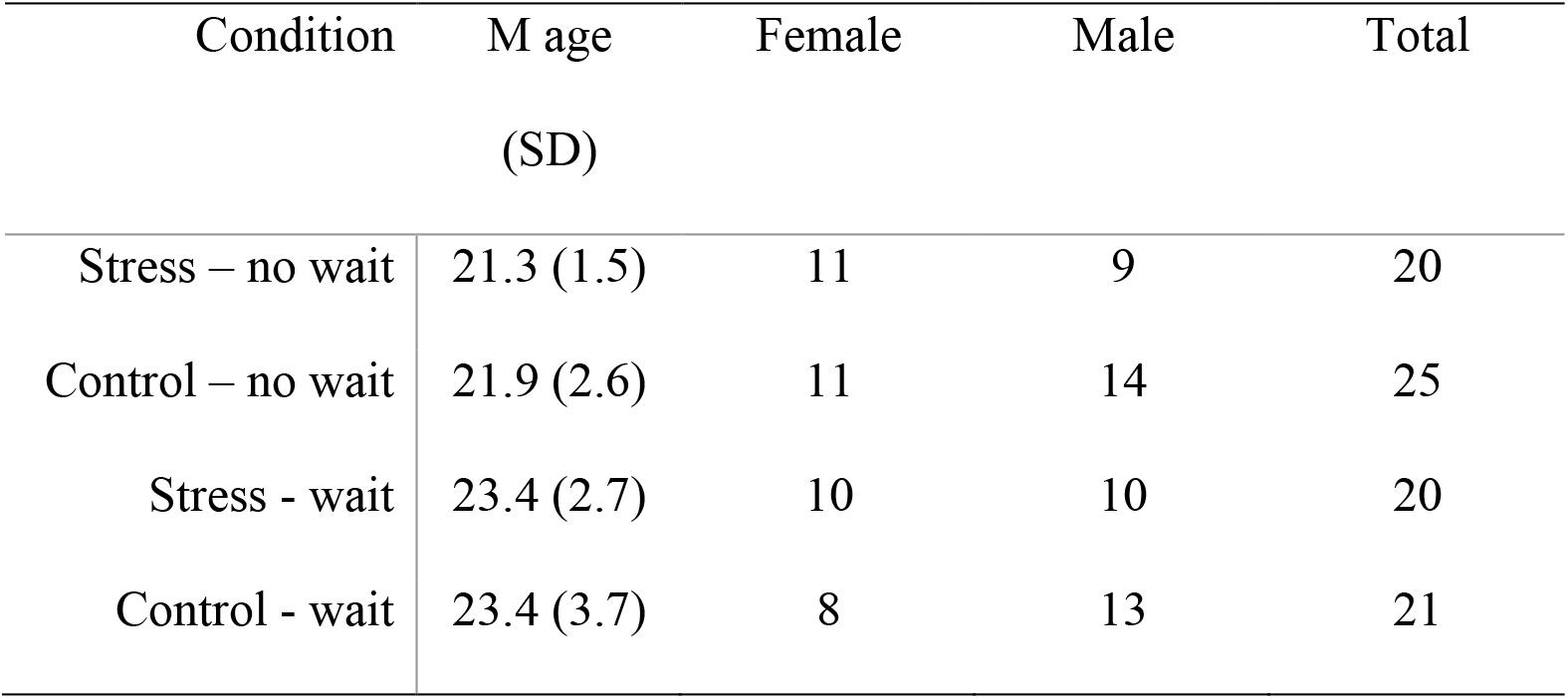
Demographic information per group.

#### Study Overview

The experiment was conducted on two consecutive days (Figure 1A), and had a 2 (stress/control) x 2 (wait/no wait) between-subject design. The first day consisted of a stress or control induction followed by a subjective stress questionnaire and an episodic learning task. The first session always took place between 12:30 and 18:00 to control for circadian cortisol rhythms. Participants either started the episodic learning task directly after the stress/control induction, or 30 minutes later, depending on the condition. Participants returned the next day for an unannounced memory test about the episodic learning task. The episodic learning task is effective in teaching participants a novel time scale and temporal structure of episodic material (Bellmund et al., 2021), and was slightly adapted. We reduced the number of trials per virtual day from 7 to 4, and administered the memory test a day later rather than shortly after learning. These changes were made to accommodate the task to the duration of the stress paradigm and to prevent potential effects of the stress induction on recall rather than just learning.

**Figure 1.**
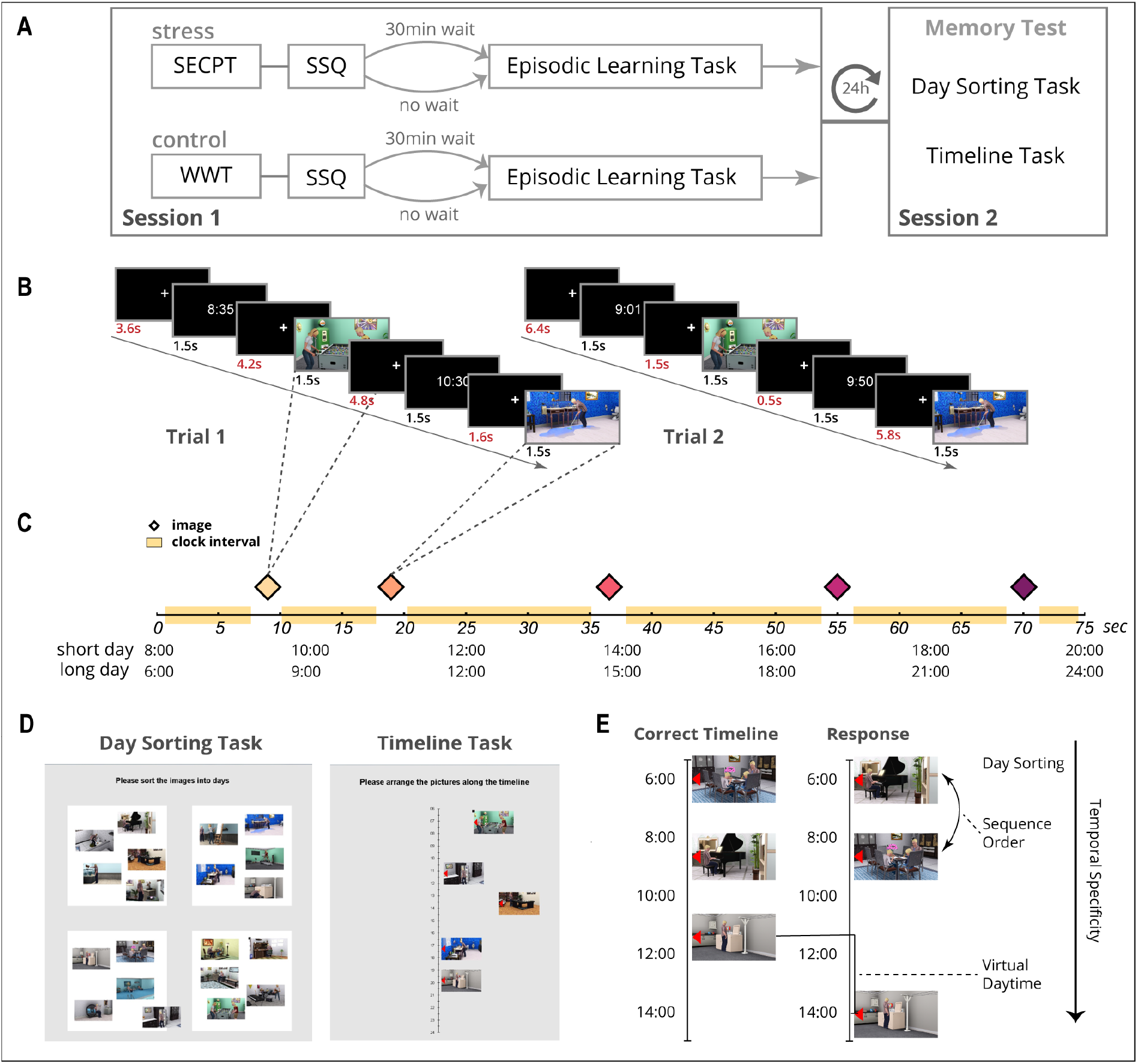
Experimental Design. **A)** Overview of study procedure for four experimental groups, over two sessions. **B)** Example of the trial structure of a virtual day, and the difference between two presentations of the same virtual day. **C)** Illustration of how the time scaling affects the virtual time distance between images for a short and long day with an identical structure. Diamonds indicate when an image is shown, and yellow blocks indicate the interval in which a clock will appear. **D)** Display of both memory tasks. **E)** Example of a response on the timeline task, and how temporal memory can be abstracted at different levels of specificity. **B-E)** The Sims 3 and screenshots of it are licensed property of Electronic Arts, Inc.

### Socially Evaluated Cold Pressor Test

#### Stress induction

The socially evaluated cold pressor test hereafter SECPT; (hereafter SECPT; Schwabe, Haddad, et al., 2008) was used to induce physiological and social stress. Participants were instructed to submerge their non-dominant hand in a bucket of ice-cold water (0-2 degrees Celsius), for a maximum duration of 3 minutes. They were told that they could remove their hand from the water if it became unbearable, but to keep it in as long as they could. As a social stressor, the test leader wore a lab coat and took on a strict and highly formal demeanor. During the task, the test leader actively monitored participants while tracking the time using a stopwatch. Additionally, participants were told that their facial expressions would be recorded with a desk camera. After 3 minutes, or earlier when participants removed their hand from the water, they were given a towel to dry their hand and were asked to complete a short subjective stress questionnaire (SSQ). The SSQ asked them to rate how stressful, painful, and unpleasant they found the task on a 10-point Likert scale (0 – 100). Participants either proceeded directly to the episodic learning task or were asked to wait 30 minutes while reading unstimulating magazines (e.g. home improvement or sailing magazines). They were not allowed to use their phone.

### Control induction

The control conditions followed the same procedure as the stress conditions. However, the SECPT was replaced by the Warm Water Test (hereafter WWT; Schwabe, Haddad, et al., 2008). The WWT has the same general structure as the SECPT, but the water is lukewarm (20-23 degrees Celsius) and none of the social stressors are applied. The test leader stayed in the room with the participant during the task to track time, but did not actively monitor participants and stayed outside of their line of sight.

### Episodic Learning Task

#### Scene images

We used 20 images, created using the life simulation video game The Sims 3 (The Sims 3 and screenshots of it are licensed property of Electronic Arts, Inc.), to construct virtual days for the episodic learning task. The images displayed everyday activities in the life of a Sim family (e.g. reading the newspaper or doing homework). All images depicted a unique scene, and they were independently rated by a sample of 40 students (Bellmund et al., 2021) as visually distinct, temporally ambiguous, and clear in terms of content. Temporal ambiguity was of particular importance as each of the 20 images was randomly assigned to one of five fixed time-points within one of 4 virtual days.

#### Virtual days

The virtual day task was designed to model the way people experience and monitor the passage of time in daily life. For example, you look at a clock which says 15:00, do some activities and afterwards estimate that it must be about 17:00 now. The virtual days represent a scaled version of this process. In the virtual days, time moves faster but clocks can be used in the same way to estimate at what time events occurred within the day, and relative to each other.

Each of the four virtual days consisted of 5 images that were randomly assigned to specific time-points (Figure 1C) within the virtual day to create a 4 unique event sequences. The time-points at which an image took place were identical for all participants, only the image assignment was randomized between participants. Participants’ task was to memorize which 5 images belonged to the same day, and to estimate at what time within the virtual day each image took place. To enable participants to estimate virtual time, a clock with the current virtual time was shown before and after each image at random intervals (Figure 1B and C). Image and clock trials were both displayed for 1.5 sec, and were interleaved by a fixation cross (Figure 1B). A 2 sec buffer between the onset of clock and image trials was used to prevent overlap due to the presentation time of the stimuli.

#### Clocks

Clocks appeared once at a random moment within specified intervals (yellow bars in Figure 1C). The time on the clocks represented the virtual time as the clock appeared on screen (see Instructional video in Supplemental Online Material), and did not change during the 1.5 sec display time. Clocks never appeared at the same moment across repetitions of the same day (Figure 1B). This allowed participants to get a closer estimate of the correct time with every repetition of a virtual day. The goal of this procedure was for participants to develop a sense of (virtual) time without relying on direct associations between clock time and an image.

#### Time Scaling

Within the virtual days, the passage of time was scaled so that each second represented a certain number of minutes in virtual time. The different time-scales allow dissociation between memory based on virtual time and real time. This feature was implemented as part of an earlier neuro-imaging study using the same task (Bellmund et al., 2021). We retained the two scales as it forced participants to learn the virtual time rather than for example counting the time in seconds. There were two short days, 8:00 – 20:00, and two long days, 6:00 – 0:00. The short and long days had the same duration in real time (75 sec) but differed in virtual time (12h and 18h). Thus, time passed more quickly in long days (1 sec = 14.4 min) than short days (1 sec = 9.6 min). In order to correctly estimate the time-point of each image, participants had to learn these time scales.

#### Episodic Learning Phase

Participants were instructed to learn the temporal structure of the 4 virtual days. They were informed that each of the images occurred at a specific time within a virtual day, and that their task was to learn which images belonged to the same day and at what time they took place using clocks. They received detailed instructions about the time scaling within the virtual days, and how they could use the clocks to estimate the time an image took place. All participants were asked to describe the task in their own words to verify that they understood the task before it started.

The episodic learning task consisted of 4 blocks of 4 event sequences (i.e. virtual days). Each virtual day was presented 4 times, once per block. An image of a moon was shown for 5 sec at the end of each virtual day as a visual boundary between the four event sequences (i.e. virtual days) in each block. The order of the four days was randomized within blocks, and there was a 30 sec break between blocks. The task took about 20 minutes to complete.

#### Memory test

One day later, participants returned to the lab for an unannounced memory test. They were told that they would do a similar task as in the first session, but the content was not disclosed. The memory test consisted of two parts (Figure 1D). In the first part, which we will refer to as the ‘day sorting task’ (Figure 1D left panel), participants were asked to recall which images belonged to the same virtual day. The 20 images appeared on the screen in a circular formation, in randomized order. Participants could drag and drop each image into one of four white squares that represented the four days. The virtual days were not given explicit labels during the episodic learning task. So participants simply grouped the five images they thought belonged to the same day in one of the squares. This task assesses the ability to recall sequence membership of the individual events (coarse level of temporal granularity).

In the second part, which we will refer to as ‘the timeline task’ (Figure 1D right panel), participants were asked to reproduce at which time each image took place within the virtual day by placing it along a timeline. The four virtual days were presented separately. A timeline was presented on screen that ran from 6:00 till 0:00, as well as the 5 images that belonged to that day. A red arrow was embedded in each image to allow more precise placement along the timeline. When participants were finished with a day they could continue with the next day. This task assesses two levels of temporal granularity, namely (from highest to lowest) memory for event sequence (sequence order) and time of occurrence (virtual event time). The memory test took about 10 to 15 minutes to complete.

### Statistical analysis

#### Subjective stress levels

To determine if the stress induction in the SECPT was successful and similar for both stress groups, we compared SSQ scores of the four experimental groups using a 2 (Condition: control, stress) x 2 (Wait: wait, no wait) between subjects ANOVA.

#### Memory test

For the day sorting task, we first tallied how many images from the same day were correctly grouped together in each square. Each square could only represent one virtual day. So if the top right square represented day 2 and contained three images of day 2, then the square received a score of 3. The scores for each day square were then added together. The maximum total score on the day sorting task was 20. This metric represents the highest level of temporal granularity (Figure 1E); clustering of events that belong to the same episode.

Performance on the timeline task was operationalized in two time metrics: sequence order and virtual daytime (Figure 1E). Scores for both metrics reflected the absolute deviation from the correct response. For both metrics, the scores were calculated per image and then averaged across all 20 images. Sequence order assessed memory accuracy for the order of the event sequence, and was operationalized as the number of positions off from the correct response (e.g. image 1 placed at position 3 produces an error score of 2). Virtual daytime assessed the ability to accurately estimate event times and create an event timeline based on the new time scales learned through the task. The virtual daytime score was calculated by subtracting the correct image time from the participants’ response (e.g. 12:40 – 12:00 = 40 virtual minutes). The time deviation was then converted from virtual minutes to seconds in real time (short days: VT / 9.6 * 60, long days: VT / 14.4 * 60) to correct for the different timescales of the long and short days (see ‘Time Scales’ above).

For both time metrics, an average score of 0 reflects perfect performance. This caused the response distribution to be skewed towards 0. Therefore, Mann-Whitney U tests were used to investigate the effect of stress induction and wait time on temporal learning. In addition to frequentist statistics, we employed Bayesian statistics to test for evidence for the null hypothesis.

##### Generalization bias

To assess the presence of a generalization bias in the virtual time estimates, we followed the analysis steps described in Bellmund et al. (2022) to produce a regression coefficient for the strength of the bias for each participant, followed by a (Condition: control, stress) x 2 (Wait: wait, no wait) between subjects ANOVA to assess group differences.

##### Exploratory analysis

To explore potential sex differences in the relationship between subjective stress and temporal memory, we conducted a linear regression in the two stress groups, with wait conducted (no wait/wait), sex (female/male) and subjective stress scores as predictors of memory performance.

## Results

### Subjective Stress

To examine whether the stress induction was successful, group differences were compared in self-reported subjective stress using a 2 (Condition: control, stress) x 2 (Wait: wait, no wait) between subjects ANOVA. We found a main effect of Condition, *F*(1, 82)= 286.14, *p* < .001, *η_p_^2^ =* .78, indicating that the SECPT indeed induced more subjective stress than the control induction (Stress: *M* = 160.8, *SD* = 59.9, Control: *M* = 11.5, *SD* = 18.6). However, there was also a significant interaction between Condition and Wait, *F*(1, 82)= 6.41, *p* = .013, *η_p_^2^ =* .073. This interaction was due to higher subjective stress scores in the Stress – Wait group (*M* = 183.0, *SD* = 62.7) compared to the Stress – No Wait group (*M* = 138.5, *SD* = 49.0), *t*(38) = -2.499, *p* = .017. This difference limits direct comparisons between the wait and no wait stress groups, and was likely due to chance given the randomized group allocation, the same protocol for both groups, and measurement of subjective stress directly after completing the SECPT. Finally, we found no sex differences in subjective stress between the stress and control group, *F*(1,82)= .5, *p* = .481, *η_p_ ^2^ =* .006, and between the stress no wait and wait group, *F*(1,36)=.016, *p* = .901,, *η_p_^2^ =* .000.

### Day Sorting Task

A Mann-Whitney U test was used to examine the effect of stress on encoding of episodic context, as indexed by which event images belong to the same virtual day. There was no significant difference in performance between the stress and control group in both the No Wait (Control: *M* (*SD*)= 12.84 (3.3); Stress: *M* (*SD*)= 13.85 (4.4)), *U* = 216.5, *p* = .442, and Wait condition (Control: *M (SD)*= 12.95 (3.4); Stress: *M (SD)*= 13.20 (3.4)), *U* = 195, *p* = .694. Furthermore, there was no difference in performance between the stress – wait and stress – no wait groups, *U* = 184.5, *p* = .678.

### Timeline Task

First, the Stress groups with their respective Control group (e.g. stress wait with control wait) were compared to examine the effect of stress on memory for temporal structure. No significant differences were found between the stress and control groups for sequence order and virtual time (No Wait: all *U* > 192, *p* > .185; Wait: all *U* > 177, *p* > .389). Second, the Stress groups were compared to examine the effect of the interval between stress and learning on memory for temporal structure. Again, no group differences were found (all *U* > 178, *p* > .552).

Bayesian independent-sample t-tests were performed in JASP (JASP Version 0.14.01) to assess evidence for the null hypothesis (i.e. no difference between the stress and control groups). A default Cauchy prior of 0.707 was used as it was difficult to determine an informed prior. Comparing the stress no wait and the control no wait group, showed anecdotal to moderate evidence for the null hypothesis across all outcome measures: Day Sorting B_10_ = .403, 95% CI [-.756, .314], Sequence order B_10_ = .307, 95% CI [-.603, .451], Virtual Daytime B_10_ = .321, 95% CI [-.641, .416]. Comparing the stress wait and the control wait group, there was again anecdotal to moderate evidence for the null hypothesis across all outcome measures: Day Sorting B_10_ = .313, 95% CI [-.605, .485], Sequence order B_10_ = .314, 95% CI [-.612, .478], Time Deviation B_10_ = .31, 95% CI [-.496, .593].

### Generalization Bias

Next, we examined whether stress before sequence learning amplifies the generalization bias reported in Bellmund et al. (2022). Replicating the analysis performed by Bellmund et al. (2022), a linear regressions for each individual participant was conducted to assess to what degree estimated event times were predicted by the average distance to other events in the same sequence position. For example, if the average virtual time of event 5 in day sequences 2-4 was relatively late compared to event 5 in day 1, did participants systematically overestimate the virtual time of the latter. To assess the presence of this bias in the individual groups, the resulting p-values from these participant-level linear models were converted to Z-scores, and tested against 0 using a permutation-based t-test. All four groups showed significant levels of this generalization bias: Control No Wait *t*(24) = 7.81, *p* = .000, *d* = 1.51, 95% CI [.98, 2.19], Control Wait *t*(20) = 7.80, *p* = .000, *d* = 1.63, 95% CI [1.04, 2.42], Stress No Wait *t*(19) = 11.82, *p* = .000, *d* = 2.53, 95% CI [1.73, 3.67], and Stress Wait *t*(19) = 5.78, *p* = .000, *d* = 1.23, 95% CI [.70, 1.93].

Then a 2 (Condition: control, stress) x 2 (Wait: wait, no wait) between subjects ANOVA was conducted on the regression coefficient (beta) of the participant-level linear models, to assess whether the generalization bias was stronger for the stress groups. There was a statistically significant interaction effect, *F*(1, 82) = 4.08, *p* = .046, *η_p_^2^ =* .047. The Stress No Wait group displayed a stronger, but not statistically significant, generalization bias than the Control No Wait group, *t*(43) = -1.7809, *p* = 0.082, *d* = .54, and Stress Wait group, *t*(38) = 1.809, *p* = 0.07, *d* = .57 (see figure 3). So, in line with Bellmund et al. (2022), all groups showed a significant generalization bias, but this bias tended to be slightly stronger (i.e. virtual time estimates were skewed more towards the average timeline) when learning took place directly after stress induction.

**Figure 2.**
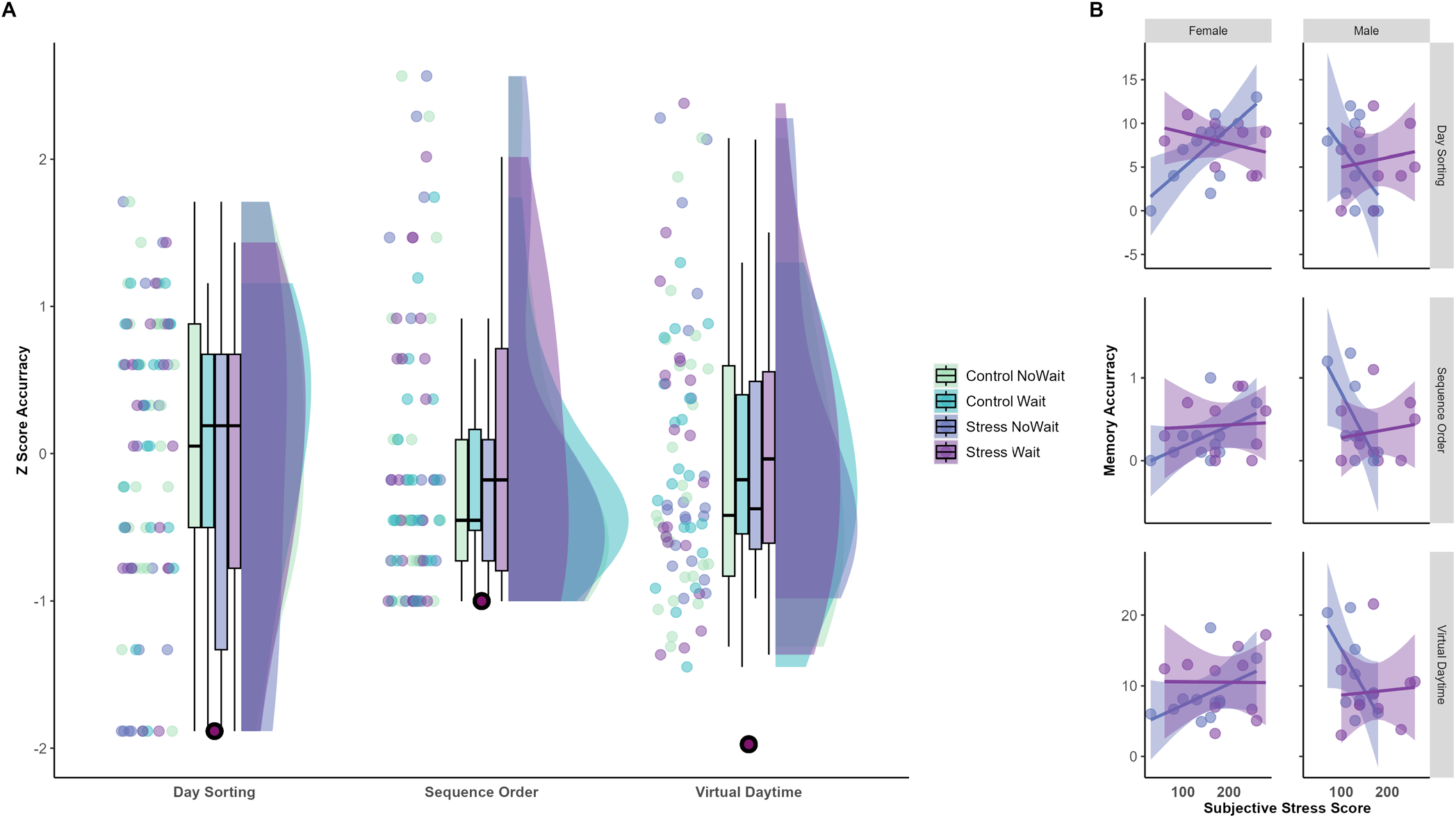
Memory performance per outcome measure per group. **A)** Data distribution of Z scores for performance on day sorting, sequence order, virtual daytime. Black dots represent the Z-score that indicates perfect performance on each measure (i.e. a score of 0). **B)** Memory accuracy as a function of subjective stress separated by sex and outcome measure, only for the stress condition. Sex interacts with subjective stress for the no wait group but not the wait group, leading to impaired memory in women when subjective stress is higher and improved memory in men when subjective stress is higher.

**Figue 3.**
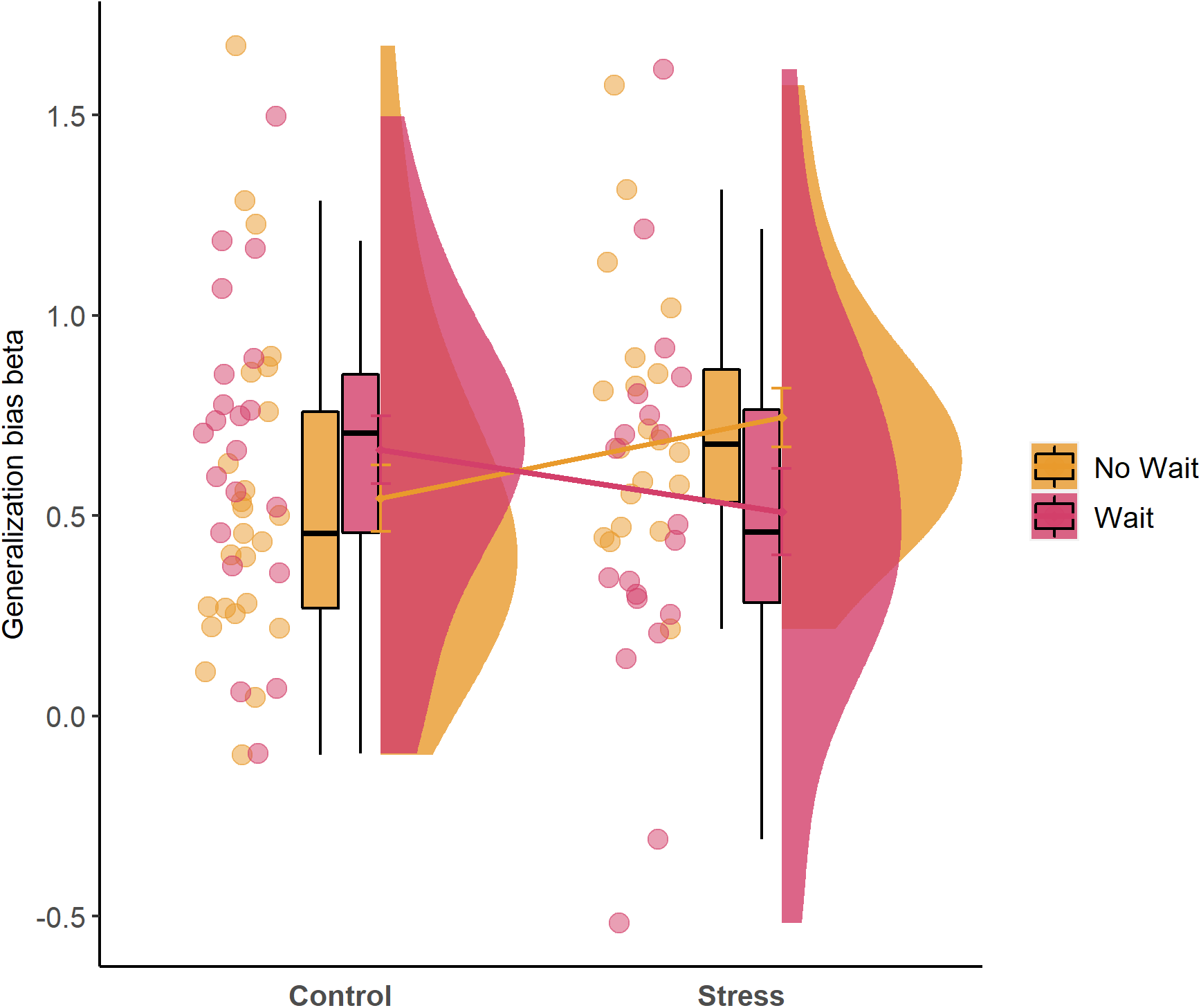
Group differences in strength of generalization bias. Higher beta indicates that participants were more likely to bias their time estimates of individual events towards a general timeline averaged across the virtual days.

### Sex differences

To explore sex differences, linear regressions were conducted to assess the interaction between wait time, sex and subjective stress on memory performance for all four time metrics in the two stress groups (wait/no wait). These analyses revealed a significant three-way interaction between wait group, sex and subjective stress predicting memory performance (see Figure 2B) for Day Sorting (*β*= .14, *p* = .012, 95% CI: .03, .24) Sequence order (*β*= .0138, *p* = .011, 95% CI: .003, .0243) and Virtual Daytime (*β*= .15, *p* = .03, 95% CI: .011, .292). Follow-up analyses, using subjective stress and sex as predictors, showed that this interaction is explained by a main effect of sex as well as the interaction between sex and subjective stress (see Supplementary table 1) in the stress no wait group (Day Sorting *F*(3,16) = 2.929, adjusted *R^2^* = .23, *p* = .06, Sequence order *F*(3,16) = 3.916, adjusted *R^2^* = .31, *p* = .02, Virtual Daytime *F*(3,16) = 3.294, adjusted *R^2^* = .26, *p* = .04), but not in the stress wait group (all *F*(3, 16) < .84, *p* > .48). The interaction pattern is quite consistent between different measures (Figure 2B).

## Discussion

This study aimed to provide a first examination of the effect of a validated stress paradigm on encoding of temporal context in episodic memories, using a fine-grained temporal memory task for episodic events sequences and event times. Following a stress or control task, participants completed an episodic learning task that has previously been proven effective in teaching participants a novel time scale and temporal structure of episodic material (Bellmund et al., 2022). While subjective stress scores indicated that the stress manipulation was successful, the results did not show a pronounced effect of stress on temporal memory.

Performance was best for the sequence order metric across groups, which seems to indicate that participants generally focused on learning the event sequence rather than the virtual time and still required a strong cue (i.e. being given the correct items per day to access that knowledge). However, we did find a slight amplification of the previously reported generalization bias effect (Bellmund et al., 2022) in the stress no wait group, compared to control no wait. Finally, our data support earlier work that found that the effect of acute stress on memory is differentially moderated by stress reactivity for men and women (Carr et al., 2016; Kluen et al., 2017). When learning took place directly after stress induction, memory accuracy improved as a function of subjective stress in men, but it deteriorated in women.

Contrary to earlier work on the effects of arousal on memory for neutral event sequences we did not find that stress enhanced (Clewett & McClay, 2021; Tambini et al., 2017), or interfered with (Huntjens et al., 2015) sequence memory. First, as suggested by Clewett et al. (2019), the enhancement of neutral sequence memory following arousal may be contingent on the neutral and arousing event sharing a task context. In the current setup, there was a clear task separation between the stress induction and the learning task. Thus, even though these tasks were performed in the same room and directly followed each other in the no wait groups, participants may have perceived a clear event boundary between the two tasks.

Second, regarding the potential negative effects of stress on sequence memory, it is possible that this effect depends on the presence of emotionally arousing or stress relevant stimuli in addition to neutral stimuli (Buchanan & Tranel, 2008; Joëls et al., 2011; van Ast et al., 2013; Wolf, 2008). The physiological stress reaction generally biases attention towards information related to the stressor, which is innately emotionally arousing (Ehlers & Todd, 2017; Sara & Bouret, 2012). This attentional bias is known to disrupt encoding of information that is deemed less relevant in the current situation in favor of information that promotes immediate survival. Previous work has shown that this attentional bias towards threat, likely mediated by noradrenergic activity, is stronger in women and leads to improved memory for negative material (Felmingham et al., 2012; Segal & Cahill, 2009). In contrast, using neutral material, our exploratory analyses showed that memory was impaired for women as a function of subjective stress. It is possible that this memory impairment is indicative of a stronger attentional bias in women that promotes memory for negative over neutral material. Future work may examine whether this effect is indeed reversed when the temporal learning task includes emotional stimuli. An exciting example of how a temporal learning task can act as the stressor is a recent study that examined temporal clustering in memory by having participants walk through a haunted house (Gregory, 2020; Reisman et al., 2021).

In line with Bellmund et al. (2022), our data show a pronounced generalization bias across all groups in the estimation of specific virtual event times. Responses systematically deviated towards the average virtual time of events that shared the same sequence position. To illustrate this effect, if you are asked which time you left for work today, and you don’t know exactly, you are more likely to guess a time that is closer to when you typically leave than one that is further away. Participants showed a stronger generalization bias for event sequences learned directly after acute stress. Previously, this generalization bias had been explained as people relying more on general knowledge to aid recollection when memory specificity is low (e.g. ‘I always leave around 8am so maybe I left around 8:05am’) (Bellmund et al., 2022).

However, this interpretation was based on a sample with high task performance, meaning that if participants lacked memory for a specific virtual event time they could indeed rely on their intact knowledge of the other event sequences. In the current study, performance across groups was quite poor which makes a ‘general knowledge’ based explanation of this bias less likely. Perhaps the stronger bias in the no wait stress group was caused by a change in learning strategy whereby participants focused on learning the general gist of the sequence structure rather than the specific event times, as participants may have been more distracted by the stress task directly prior. This might similarly lead to a generalized timeline in memory for virtual event times, as all sequences are encoded following the same gist-like template.

This interpretation, i.e. following acute stress participants rely on simpler heuristics to solve a task, corresponds with findings that in high pressure situations participants are more likely to rely on simpler problem-solving strategies thereby reducing their performance accuracy (Beilock & Decaro, 2007; Plessow et al., 2012). Evidence from work on math proficiency has shown that stress and time pressure impair performance by overloading working memory (Caviola et al., 2017; Plessow et al., 2012). Demanding computations, such as those required to deduce virtual time in this task, are more difficult to achieve when working memory resources are low and can in and of themselves introduce pressure (Lemaire & Callies, 2009). Finally, stress affects working memory performance differentially in men and women. Similar to our findings, research has shown that acute stress enhances response time on an n-back working memory task men and deteriorates response time in women (Schoofs et al., 2013). While we did not directly manipulate or assess working memory in the current experiment, the generalization bias pattern and sex interaction following acute stress are in line with a working-memory based account.

Furthermore, our data suggest that the level of subjective stress modulates the effect of acute stress on memory, and that it does so differently for men and women. Specifically, acute stress impaired memory in women and enhanced memory in men as a function of subjective stress. While our sample size does not allow us to draw stark conclusions, this finding does mirror previous work on sex differences in noradrenalin activity (Carr et al., 2016) and its relation to memory performance. When yohimbine, a noradrenalin antagonist, was administered prior to learning, performance on a memory generalization task with neutral stimuli was impaired in women and improved slightly in men (Kluen et al., 2017).

Speculatively, this could suggest that stress reactivity, both physiological and subjective, differentially affects learning in men and women but this modulation does not appear to be specific to a particular type of memory nor did it induce a discernible shift in learning strategy.

A limitation of the current study is that both the control and stress groups had high error scores on the virtual time measure (mean error of about 1.5 – 2.5 virtual hours) compared to earlier work using the same task (Bellmund et al., 2022). This difference in performance may be due to changes in the timing of the memory test, which was conducted a day later in the current experiment instead of directly after learning (Bellmund et al., 2022). The reason for this change was to limit the effects of the stress task on learning. If the memory test was performed directly after learning, stress might also interfere with recollection (Elzinga et al., 2005). This larger delay between learning and recall may have affected the accuracy of temporal memory across groups, which may have obscured stress-related effects. Future work using this paradigm could increase the number of learning trials to increase sensitivity of the task at delayed recall.

A further limitation is the lack of an objective measure of stress reactivity. Both subjective stress and cortisol responses are known to vary widely between people due to factors like sex, age, menstrual cycle and chronic stress (Kirschbaum et al., 1999; Kudielka et al., 2009). In addition, subjective stress and physiological stress reactivity do not always correlate well (Campbell & Ehlert, 2012; Duchesne & Pruessner, 2013). Therefore, future work should consider including both salivary cortisol as measures of stress reactivity next to subjective stress measurements. Indeed, one study demonstrated that endogenous cortisol secretion moderated the impairing effect of stress on implicit spatial learning (Meyer et al., 2011).

In summary, this study examined the effect of stress on encoding of temporal context information in episodic memory. The results do not show a discernible impact of stress on the ability to remember temporal context, across levels of temporal granularity. We did observe opposing effects of acute stress on general memory ability for men and women depending on the level of subjective stress. Future work should consider including tasks that are more sensitive to subtle changes in temporal context memory and measures of physiological stress reactivity to further disentangle this interaction between sex, (subjective) stress reactivity and memory performance.

## Acknowledgements

We would like to thank Christian Doeller and Lorena Deuker for permitting us to use the episodic learning task that was originally developed by Lorena Deuker and NDM as part of Bellmund et al. (2021). We thank Maja Kalkofen and Vanessa Danzer for their assistance with data collection.

Open Practices Statement. The experiment reported in this article was not preregistered. Requests for raw data and materials can be e-mailed to the corresponding author. Pre- processed data that were used for the main analyses can be found at: https://osf.io/85g3x/?view_only=6171b06689924bf6aba88a1d393ba1aa.

## Declarations

### Ethics approval

The study was approved by the institutional ethical review board at Utrecht University (FETC16-090).

### Consent to participate

Informed consent was obtained from all individual participants included in the study.

### Consent for publication

The authors affirm that human research participants provided informed consent for publication.

### Availability of data and materials

The datasets generated during and/or analyzed during the current study can be found at https://osf.io/85g3x/?view_only=6171b06689924bf6aba88a1d393ba1aa. Materials and raw data are available from the corresponding author on request.

### Code availability

Not applicable.

### Author Contributions

NDM, LG and IME designed the study; NDM collected the data; NDM and LG analyzed the data; all authors interpreted the data; NDM wrote the first draft of the article, and LG and IME provided critical revisions. All authors approved the final version of the manuscript for submission.

